# Thermal constraints shape metabolic scaling and distribution of the tufted ghost crab (*Ocypode cursor*) in the Mediterranean Sea

**DOI:** 10.64898/2026.01.14.699443

**Authors:** Vojsava Gjoni, Guillaume Marchessaux, Mario F. Tantillo, Gianluca Sarà

## Abstract

The Mediterranean Sea is undergoing significant environmental changes due to climate change and the introduction of non-native species, impacting biodiversity and ecosystem dynamics. *Ocypode cursor* (tufted ghost crab) has expanded its range, likely in response to changing thermal conditions.This study investigates the metabolic scaling of *O. cursor* across three active months (July, August, and September) under air- and water-breathing conditions during both daytime and nighttime. The results revealed seasonal variation in metabolic scaling, with significant differences in intercept values and scaling slopes among months. During daytime air-breathing conditions, metabolic rates increased in August and September regardless of body size, while at night in August, larger crabs exhibited higher metabolic rates. Under water-breathing conditions, smaller crabs showed greater metabolic responses in August during the day, whereas nighttime activity remained stable across months. These results indicate that temperature, diel cycle, and respiratory mode shape metabolic scaling as a comparative indicator of thermal performance rather than a direct proxy for fitness. Understanding these responses provides insight into *O. cursor*’s ecological flexibility and contributes to assessing ectotherm responses to ongoing Mediterranean warming.

## INTRODUCTION

The Mediterranean Sea faces well-documented environmental challenges, including the introduction of non-native species, all of which threaten its biodiversity (Coll et al., 2010). Climate change compounds these issues, with rising sea temperatures and acidification impacting the distribution and abundance of marine species (Findlay & Turley, 2021; Hastings et al., 2020). Over the past century, the Mediterranean has warmed by approximately 1.5°C, leading to species shifts toward cooler waters or deeper depths, and population declines in others (Pastor et al., 2019; Free et al., 2020). These shifts disrupt community structures, with species expanding into previously unsuitable areas, altering ecological balances (Freeman et al., 2018; Poloczanska et al., 2016). The tufted ghost crab (*Ocypode cursor*), native to the Atlantic Ocean and southeastern Mediterranean, exemplifies these concerns. Since the 1980s, this species has expanded its range, likely due to warming seas (Marchessaux et al., 2024). Inhabiting sandy beaches and dunes, these small, agile crabs are ecologically significant as both predators and scavengers, feeding on mollusks, crustaceans, and carrion while also serving as prey for birds and fish (Schuchman & Warburg, 1978; Schlacher & Lucrezi, 2014).

Temperature is a key factor influencing physiological and ecological processes across organisms (Brown et al., 2004; McCain and King 2014). Consequently, research on non-native species has focused on how temperature impacts critical traits like body size and metabolic rate (Brown et al., 2004; Glazier 2015; Lindmark et al., 2004; O’Connor 2009).

According to the Metabolic Theory of Ecology (MTE), metabolic rates increase exponentially with temperature, independent of body size, suggesting a constant metabolic scaling slope across temperature gradients. However, deviations from this universal slope have been reported, indicating size-dependent metabolic responses to temperature changes (Glazier 2014; Glazier 2018; Glazier 2020; Killen et al., 2010; Glazier 2005). These deviations align with the metabolic-level boundaries hypothesis (MLBH), which proposes that rising temperatures amplify surface-area-related metabolic processes relative to volume-related ones, shifting metabolic scaling slopes from 1 to 2/3 in isomorphic organisms (Glazier 2014; Killen et al., 2010; Glazier 2005). Despite these insights, limited research exists on the metabolic scaling of non-indigenous species (NIS) and how environmental factors shape it. Temperature is known to exert immediate phenotypic effects on the mass-specific metabolic rate of certain species (Glazier 2014; Glazier 2020; Killen et al., 2010; White & Kearney 2011. However, the effect of temperature on metabolic scaling are underexplored in non-native species, leaving significant gaps in understanding their physiological and ecological adaptations.

*O. cursor* is well-known for its burrowing behavior, constructing deep sand cavities that can extend to depths of up to 1 meter (Schuchman & Warburg, 1978; Shiber & Izzidin, 1978; Strachan et al., 2016; Tiralongo et al., 2020). These burrows serve as vital refuges, offering protection from both predators and extreme environmental conditions, such as intense summer heat, which can reach up to 50°C in areas like Greece (Strachan et al., 2016). The burrows maintain a cooler and more humid microclimate, which is essential for the crab’s respiration during the hottest parts of the day (Chan et al., 2006). Structurally, the burrows are elongated and tunnel-shaped, often featuring an adjoining chamber where the crab can retreat for safety and protection from external disturbances (Schuchman & Warburg, 1978; Shiber & Izzidin, 1978). In adittion, *O. cursor* exhibits a diverse feeding strategy, acting as a scavenger and predator. It feeds on animal carcasses, plant organic matter, and human food remnants, such as picnic leftovers on beaches (Strachan et al., 2016). Additionally, it preys on marine invertebrates and is frequently observed foraging at night near or along the water’s edge, where it can access a broader range of food sources (Strachan et al., 2016).

Here, we measured the metabolic scaling with body size of *O. cursor* during its three primary active months: July, August, and September. Measurements were conducted during both daytime and nighttime to account for the organism’s behavioral changes across different activity periods. This dual approach allowed us to capture variations in metabolic scaling associated with the diurnal and nocturnal activity patterns of *O. cursor*. Additionally, we assessed metabolic scaling under both air-breathing and water-breathing conditions, reflecting the natural environments this species encounters. By comparing metabolic rates in these two conditions, we were able to evaluate how respiration in relation with body size vary based on the medium the crabs are exposed to, as well as how these factors shift throughout the active months. This comprehensive approach enabled us to test the behavioral and physiological changes in *O. cursor* across different temporal and environmental contexts.

Specifically, we explored how metabolic scaling responds to seasonal changes, diel activity patterns, and the transition between air and water conditions. These insights provide a better understanding of the species’ adaptability and energy strategies, shedding light on its ecological role and how it manages physiological demands in response to environmental variability.

This species exploit these changes by adapting their diets, migration, or reproduction to synchronize with shifting seasonal cycles (Langan et al., 2021). Their burrows provide shelter and regulate temperature, supporting their survival and potentially altering the physical and ecological characteristics of invaded areas (Yilmaz & Barlas, 2020). This range expansion raises critical questions about their role in influencing biodiversity and contributing to habitat changes, such as beach erosion, in regions where they were previously absent. While these adaptive strategies may offer short-term survival advantages, they often pose threats to native species by modifying niches and ecosystem dynamics. Understanding these adaptations is critical for conservation strategies, though it is acknowledged that such responses may not sustain long-term resilience, particularly for specialists and stenotherms highly vulnerable to climate change (Brakes et al., 2021).

## MATERIALS AND METHODS

### Measure of metabolism

The study was carried out during three months of activity: July, August and September 2024; under air-breathing and water-breathing conditions, day and night.

Specimens were captured during dusk on the square in Menfi (Sicily, GPS) using a handnet, and individuals were individually placed in tubs (5 L volume) containing sand at the bottom to acclimatize them before experiments. For each experiment, crabs of different sizes were used to explore the MTE. Experiments were carried out directly on the beach in the ghost crab habitat. Measurements were taken at 20.00 (early evening) and 06.00 (before sunrise). The experience where replicated 3 times on 3 days with different specimens.

A breathing chamber system was developed to measure ghost crab metabolism directly in the field (Figure 1). The breathing chamber consisted of a circular container with a volume of 2,300 mL, the lid of which was placed on the sand. Inside the breathing chamber, a ventilation system, powered by a battery positioned outside the chamber, homogenized the air (Marchessaux et al., 2024). A membrane was also bonded to the inside of the chamber, enabling oxygen consumption to be measured using optical fibers.

**Figure 1.**
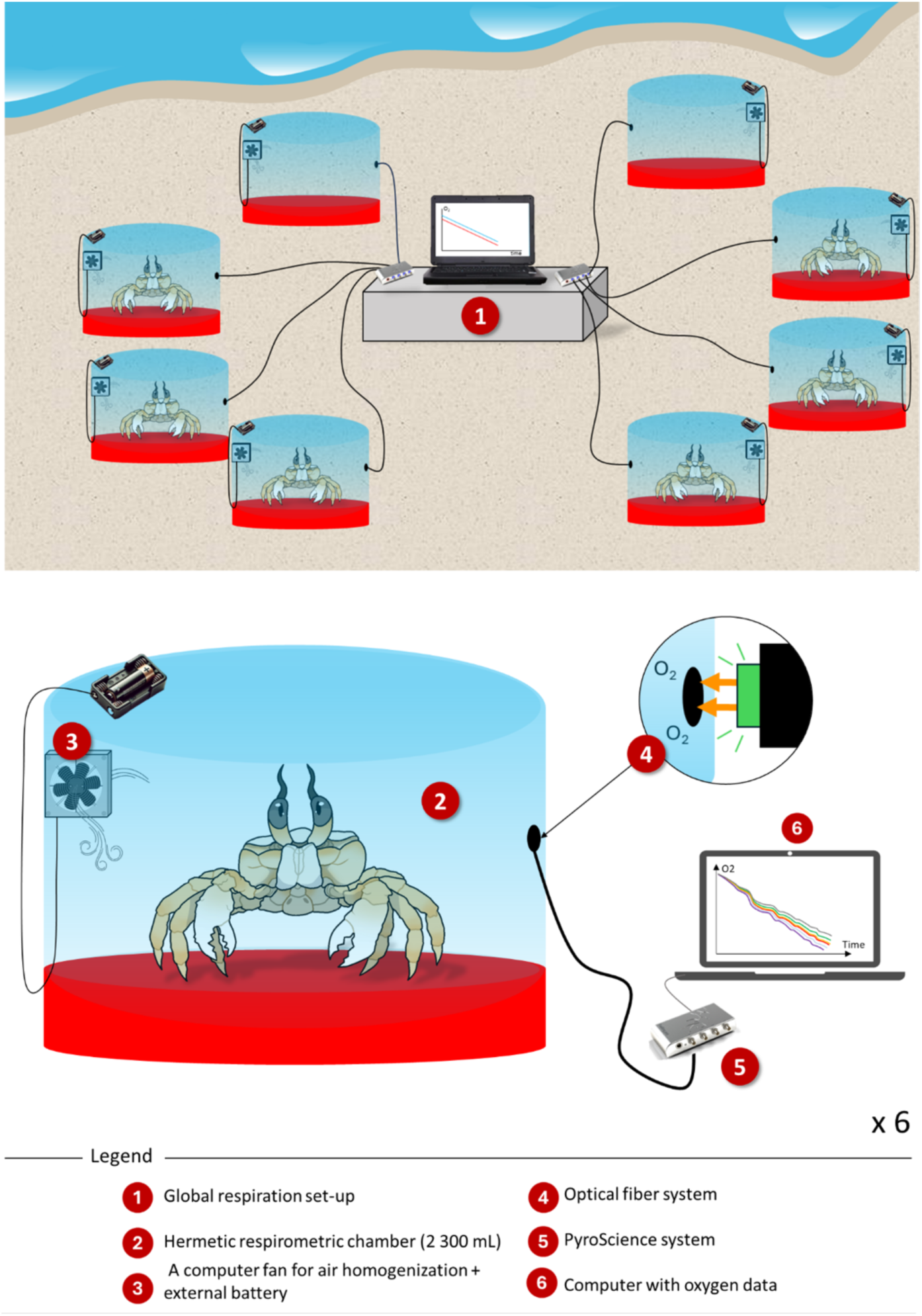
Experimental setup for measuring whole-organism respiration in crabs. A computer-based monitoring system (1) simultaneously records oxygen consumption from six hermetically sealed respirometric chambers (2), each containing a single crab. Inside each chamber, a small battery-powered fan (3) ensures air homogenization. Oxygen concentration is measured using an optical fiber sensor (4) connected to a PyroScience optode system (5). Data are logged continuously on a computer (6), allowing the quantification of oxygen decline over time.

Respiration measurements of the specimens were carried out on the same individuals in air for a minimum of 1 hour, then in seawater (at least 1 hour of measurement) to determine the change in their metabolism based on the habitat in which ghost crabs live. Respiration rates (RR, mgO2 h-1 gWW-1) were measured continuously for 1 hour using the Pyro Science Firesting O2 system.

Six *Ocypode cursor* specimens were used, placed individually in a respiration chamber. Two other empty chambers were used as controls. Respiration rate was calculated using the formula (Sarà et al. 2013): 𝑅𝑅 = (𝐶_𝑡0_ − 𝐶_𝑡1_) 𝑉𝑜𝑙_𝑟_ 60 (𝑡_1_ − 𝑡_0_)^−1^, where Ct_0_ is the initial oxygen concentration, Ct_1_ the final oxygen concentration, and Vol_r_ the chamber’s volume. At the end of the experiment, biometric parameters including carapace width (CW) and wet weight (WW) were measured to adjust respiration rates according to individual wet weight, taking into account the effects of body mass.

### Statistical analysis

We quantified the metabolic scaling of *Ocypode cursor* and its dependence on breathing medium (air versus water), time of day (day versus night), and body size using linear models. In line with the Metabolic Theory of Ecology (MTE), we formulated *a priori* hypotheses predicting (i) a positive allometric relationship between respiration rate and body mass, and (ii) potential shifts in the scaling slope and intercept associated with environmental conditions (breathing medium and diel period).

Respiration rate (RR; mg O₂ h⁻¹ g WW⁻¹) was used as a proxy for metabolic rate. To evaluate metabolic scaling, both RR and wet weight (WW) were log₁₀-transformed prior to analyses. Log-transformation linearizes power-law relationships and stabilizes variance. All continuous predictors were centred and standardized to improve model convergence and interpretability.

We used Bayesian inference to estimate model parameters, as this approach allows flexible modelling of hierarchical data structures and robust estimation of uncertainty. The response variable was assumed to follow a normal distribution:

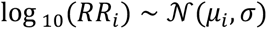

with the linear predictor defined as:

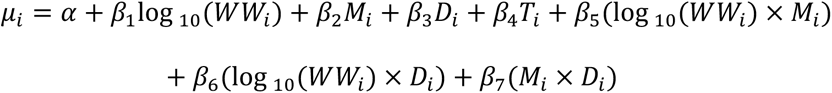

where 𝛼 is the intercept, 𝛽_1_represents the metabolic scaling slope with body mass, 𝑀_𝑖_ is the breathing medium (air vs water), 𝐷_𝑖_ is the diel period (day vs night), and 𝑇_𝑖_ represents the experimental replicate (sampling day). Interaction terms were included to test whether metabolic scaling differed among environmental conditions. Individual crabs were treated as independent observations, as different specimens were used in each experimental replicate.

Following MTE expectations, the prior distribution for the body-mass scaling coefficient was specified as a normal distribution centred on 0.75 (Normal(0.75, 2)), reflecting the predicted scaling exponent while remaining weakly informative. All other fixed effects were assigned Normal(0, 1) priors, and the residual standard deviation (𝜎) was assigned an Exponential(1) prior.

Models were fitted using Markov chain Monte Carlo (MCMC) sampling with the No-U-Turn Sampler implemented in *Stan*, accessed via the *brms* package (Bürkner 2017). Four chains were run for 4,000 iterations each, with the first half discarded as warm-up. Convergence was assessed using the Gelman–Rubin statistic (R^), ensuring all values were < 1.1 (Gelman & Rubin 1992), and by visual inspection of trace plots. Model adequacy was evaluated using posterior predictive checks (Conn et al. 2018; Hooten & Hobbs 2015).

Results are reported as posterior means with 95% credible intervals (CrI). Differences among treatments were interpreted as biologically meaningful when posterior distributions showed limited overlap and when the probability of directional effects exceeded 95%.

## RESULTS

The ghost crab (*O. cursor*) exhibited significant changes in the scaling of metabolic rate with body mass over its three active months (July, August, and September), influenced by the type of respiration (air or water). These shifts varied between daytime and nighttime, reflecting how *O. cursor* adjusts its metabolic processes in response to environmental conditions and activity cycles. During the day, metabolic rates under both air and water conditions showed distinct patterns compared to nighttime. This variation underscores the crab’s capacity for physiological adaptation to environmental factors such as temperature and oxygen availability.

Under air-breathing conditions during the daytime, the metabolic scaling of ghost crabs revealed seasonal differences in intercept values across months, while the scaling slopes remained largely unchanged. Specifically, the intercepts were higher in August and September compared to July, shifting from −0.13 in July to 0.23 in August and −0.31 to 0.24 in September (Table 1). The probability of observing a higher intercept was 0.83% in August and 0.75% in September relative to July. Meanwhile, the probabilities for steeper metabolic scaling slopes were 0.05% in August and 0.09% in September. These results suggest that, in both August and September, metabolic rates increased regardless of body size when compared to July. Under air-breathing conditions during the nighttime, ghost crabs showed different metabolic responses, with only August showing higher intercepts and steeper metabolic scaling slope compared to other months (Figure 1, Table 1). The probability of observing a higher intercept in August was 0.05%, while in September it was 0.08%, both relative to July. For steeper metabolic scaling slopes, the probabilities were 0.85% in August, 0.09% in July, and 0.06% in September. This suggests that during August nights, larger crabs increased their metabolic rates more significantly than smaller individuals.

Under water-breathing conditions during the daytime, ghost crabs exhibited distinct metabolic patterns, with August showing a lower metabolic scaling slope compared to July and September (Figure 2, Table 1). The probability of a higher intercept in August was 0.05%, increasing slightly to 0.08% in September relative to July. For lower scaling slopes, the probabilities were 0.85% in August, 0.09% in July, and 0.06% in September. These findings indicate that in August, smaller crabs had a more pronounced metabolic rate increase compared to larger individuals. During nighttime water-breathing conditions, metabolic scaling showed no significant seasonal variations in intercept or slope across the three months. The probability of a higher intercept was 0.03% in August and 0.05% in September relative to July. For steeper metabolic scaling slopes, the probabilities were 0.05% in August and 0.09% in September. These results suggest that under nighttime water-breathing conditions, the metabolic responses of ghost crabs remained consistent throughout the active months.

**Figure 2.**
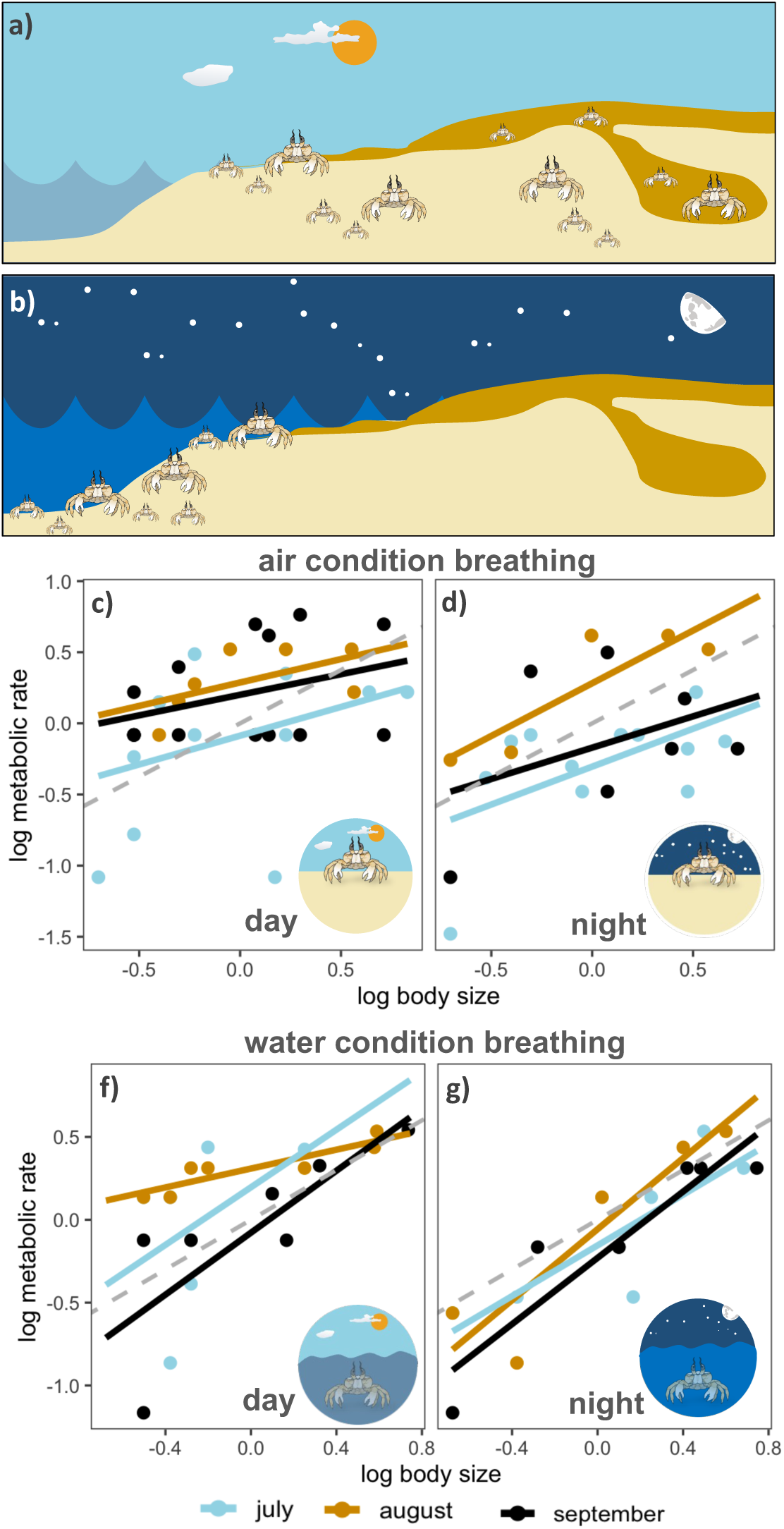
**(a)** *O. cursor* constructs deep sand burrows, up to 1 m, offering protection from predators and extreme heat. These burrows maintain cool, humid conditions that facilitate breathing during the hottest hours of the day. **(b)** At night, *O. cursor* becomes active near or in the water, scavenging for carcasses, plant material, and marine invertebrates. (**c-f)** Relationships between metabolic rate and wet body mass of *O. cursor* populations across active months (July, August, September): **(c)** Air-breathing during the day, (**d)** Air-breathing during the night, **(e)** Water-breathing during the day, and (**f)** Water-breathing during the night. Grey dashed lines represent the expected 0.75 slope predicted by the Metabolic Theory of Ecology (MTE).

**Figure 3.**
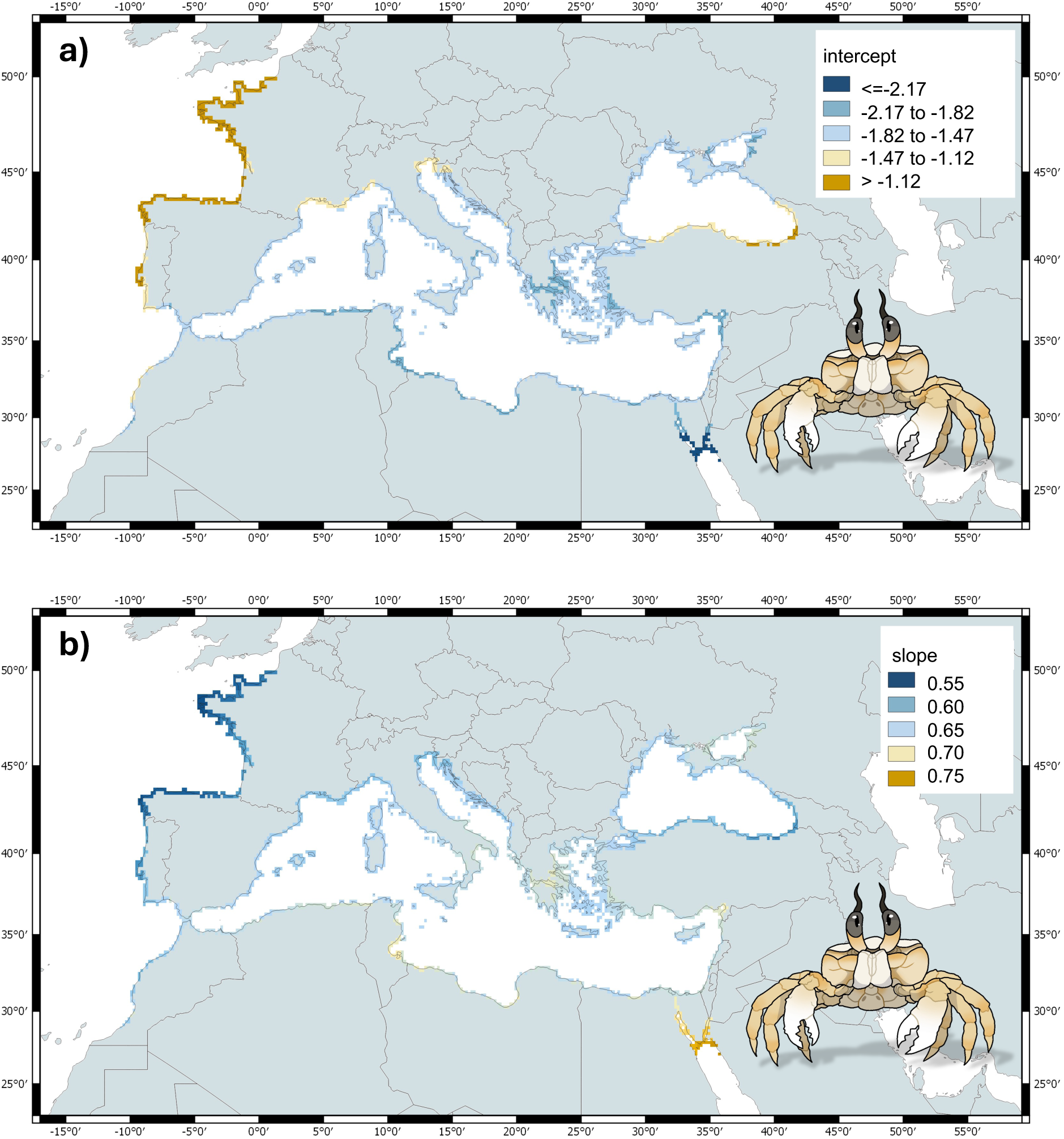
Spatial variation in predicted metabolic scaling parameters of the ghost crab *Ocypode cursor* across the Mediterranean Sea and adjacent regions. **(a)** Intercept values, representing baseline metabolic rates independent of body size. Higher intercepts (yellow) indicate areas where all individuals, regardless of size, exhibit elevated metabolic activity. **(b)** Slope values, representing the scaling relationship between body mass and metabolism. Steeper slopes (yellow) highlight regions where larger crabs experience disproportionately higher metabolic rates, while shallower slopes (blue) indicate stronger metabolic responses in smaller individuals. Together, these maps illustrate spatial heterogeneity in the energetic strategies of *O. cursor* populations, with implications for their resilience and ecological impact under ongoing climate change.

## DISCUSSION

Our study demonstrates how the metabolic scaling of Ocypode cursor varies with season, diel cycle, and respiration mode, underscoring the complex interplay between body size, activity patterns, and environmental drivers in shaping energetic strategies. These findings refine our understanding of ghost crab physiology within Mediterranean coastal systems, which are currently undergoing rapid warming and increasing anthropogenic pressure (Coll et al., 2010; Pastor et al., 2019; Marchessaux et al., 2024). By integrating metabolic scaling theory with field-based respiration measurements, our results contribute a mechanistic perspective on how physiological performance may mediate the ongoing redistribution of thermophilic species across the Mediterranean basin.

By quantifying metabolic scaling across multiple temporal and respiratory contexts, we reveal a marked physiological plasticity that allows O. cursor to adjust its energy dynamics in response to fluctuating thermal and environmental conditions. This plasticity is consistent with a broader ecological strategy in which burrow use, nocturnal foraging, and bimodal respiration act as complementary buffering mechanisms against thermal and desiccation stress (Lucrezi & Schlacher, 2014; Strachan et al., 2016). Such behavioral– physiological coupling is particularly relevant in sandy beach ecosystems, where operative temperatures can deviate substantially from ambient air temperatures and exceed physiological limits during summer daytime conditions (Sunday et al., 2014).

Importantly, our physiological findings should be interpreted with caution with respect to ecological outcomes. Metrics such as thermal performance breadth, thermal optimum (Topt), or thermal habitat suitability (THS) do not directly quantify fitness, survival, or population growth. Rather, they represent physiological indices of thermal adequacy, useful for comparative analyses across seasons, habitats, or geographic regions. As emphasized in broader macrophysiological frameworks, physiological tolerance alone rarely predicts persistence without considering behavioral thermoregulation, ecological interactions, and habitat availability (Huey et al., 2012; Gunderson & Stillman, 2015). In this sense, THS should be viewed as a relative indicator of thermal suitability, not a direct proxy for demographic performance.

Daytime intercept values increased in August and September, indicating elevated baseline metabolic rates across all body sizes. This pattern suggests that midsummer conditions impose higher energetic costs irrespective of individual mass, likely reflecting increased maintenance demands under thermal stress (Brown et al., 2004; Glazier, 2018). Empirical measurements from eastern Mediterranean beaches show that surface sand temperatures frequently exceed 50 °C at midday, while burrow microclimates remain comparatively cooler and more stable (Strachan et al., 2016). Elevated intercepts may therefore reflect the energetic costs associated with maintaining physiological homeostasis during brief surface activity or sustaining burrow ventilation under extreme heat. The absence of changes in scaling slopes indicates that this stress acts uniformly across size classes rather than selectively on larger or smaller individuals.

In contrast, nighttime conditions in August were characterized by increases in both intercepts and slopes, with larger individuals exhibiting disproportionately higher metabolic rates. This pattern aligns with the metabolic-level boundaries hypothesis, which predicts shifts in scaling relationships under heightened energetic demand (Glazier, 2014; Killen et al., 2010). Larger crabs may intensify nocturnal activities—such as long-distance foraging, burrow maintenance, or reproductive behavior—during peak summer months. Observational studies have documented that adult Ocypode individuals range farther up the beach at night and actively prey on turtle hatchlings and bivalves (Strachan et al., 1999, 2016), behaviors likely associated with increased metabolic expenditure. Our results thus provide physiological support for previously observed behavioral patterns, linking diel activity rhythms to size-dependent energetic costs.

Under daytime water-breathing conditions in August, smaller crabs exhibited greater metabolic responses than larger individuals. Juveniles are likely more constrained by respiratory inefficiency, higher surface-area-to-volume ratios, and shallower burrow use (Shuchman & Warburg, 1978; Rodrigues et al., 2016). This ontogenetic sensitivity is particularly relevant in the context of climate warming, as theoretical and empirical studies indicate that ectotherms often experience reduced thermal safety margins at early life stages (Sinclair et al., 2016; Sunday et al., 2014). Larger individuals, by contrast, may benefit from deeper burrows and more effective air-breathing, reducing their dependence on energetically costly aquatic respiration.

Nighttime water-breathing showed stable metabolic scaling across months, suggesting that aquatic nocturnal excursions impose consistent energetic demands regardless of seasonal temperature variation. Ghost crabs display highly predictable nocturnal foraging patterns, with activity peaks at dusk and dawn (Strachan et al., 1999; Lucrezi & Schlacher, 2014). Such consistency may enhance resilience by ensuring reliable energy acquisition even during periods of elevated thermal stress.

Taken together, these results highlight a context-dependent modulation of metabolic strategies in O. cursor. Daytime thermal stress elevates baseline metabolic costs, nocturnal activity disproportionately affects larger individuals, and juvenile crabs experience greater vulnerability during aquatic respiration. These findings reinforce the notion that thermal performance is shaped not only by mean temperatures, but also by diel variability, behavioral buffering, and life-history stage (Gunderson & Stillman, 2015).

Within this framework, the concept of a “thermal corridor” warrants clearer contextualization. Rather than implying a simple or deterministic expansion pathway, a thermal corridor should be understood as a spatio-temporal window of physiological permissiveness, where warming relaxes cold constraints while behavioral mechanisms—such as burrow use and nocturnality—buffer against acute heat stress. Similar concepts have been invoked in macrophysiological and biogeographic studies, where species persistence depends on the interaction between physiological limits, behavioral thermoregulation, and landscape connectivity (Huey et al., 2012; Sunday et al., 2014).

Crucially, linking thermal optima and THS to regional climate projections enhances the predictive value of this framework. Mediterranean summer temperatures are projected to increase by approximately +2–3 °C over the coming decades (Pastor et al., 2019), a magnitude comparable to shifts shown to substantially erode thermal safety margins in ectotherms (Gunderson & Stillman, 2015). Under such scenarios, behavioral buffering may allow O. cursor to exploit newly suitable northern habitats while maintaining performance within existing ranges, at least until upper thermal limits are approached. This interpretation aligns with broader evidence that distributional shifts in marine and coastal ectotherms are often mediated by physiological tolerances interacting with behaviorally accessible refugia (Stuart-Smith et al., 2015).

## CONCLUSION

The Mediterranean Sea is among the fastest warming marine regions globally (Pastor et al., 2019; Free et al., 2020). The capacity of Ocypode cursor to modulate metabolic scaling across seasons, diel cycles, and respiratory modes likely underpins its recent expansion into central and western Mediterranean shorelines (Deidun et al., 2017; Marchessaux et al., 2024). However, physiological flexibility should not be equated with unlimited resilience. Indices such as THS and Topt provide valuable comparative tools for assessing thermal suitability across space and time, but they do not directly predict fitness, survival, or population growth.

As both predator and scavenger, O. cursor plays a significant role in sandy beach food webs and is increasingly recognized as a bioindicator of beach condition (Barros, 2001; Lucrezi, 2015). Our results add a physiological dimension to this role by identifying size- and context-dependent energetic constraints that may shape population responses to future warming. Populations dominated by large individuals may face rising energetic costs during nocturnal activity, while juvenile-dominated populations at expanding range margins may be particularly sensitive to aquatic respiration stress.

By integrating metabolic scaling, behavioral buffering, and climate projections, our study supports a nuanced view of ghost crab expansion along Mediterranean coasts: not as a simple poleward shift, but as a progression along a thermally permissive corridor shaped by physiological limits, behavioral plasticity, and habitat structure. Incorporating these mechanisms into predictive frameworks will be essential for anticipating the ecological consequences of ongoing coastal warming and managing sandy beach ecosystems under future climate scenarios.

